# Structure of the TXNL1-bound proteasome

**DOI:** 10.1101/2024.11.08.622741

**Authors:** Jingjing Gao, Christopher Nardone, Matthew C.J. Yip, Haruka Chino, Xin Gu, Zachary Mirman, Michael J. Rale, Joao A. Paulo, Stephen J. Elledge, Sichen Shao

**Affiliations:** Department of Cell Biology, Harvard Medical School, Boston, MA 02115; Department of Genetics, Harvard Medical School, Boston, MA 02115; Division of Genetics, Department of Medicine, Brigham and Women’s Hospital, Boston, MA 02115; Department of Neurobiology, Harvard Medical School, Boston, MA 02115; Howard Hughes Medical Institute, Boston, MA 02115

## Abstract

Proteasomes degrade diverse proteins in different cellular contexts through incompletely defined regulatory mechanisms. Here, we report the cryo-EM structure of thioredoxin-like protein 1 (TXNL1) bound to the 19S regulatory particle of proteasomes via interactions with PSMD1/Rpn2, PSMD4/Rpn10, and PSMD14/Rpn11. These interactions are necessary for the ubiquitin-independent degradation of TXNL1 upon cellular exposure to metal- or metalloid-containing oxidative agents, thereby establishing a structural basis for the stress-induced degradation of TXNL1.

---

Proteasomes are the primary protein degradation machinery in eukaryotes and comprise a 20S core particle harboring proteases capped by one or two 19S regulatory particles (RPs)^1^. Conventionally, proteasomes degrade poly-ubiquitylated proteins that are recruited by ubiquitin receptors on the 19S RP: PSMD2/Rpn1, PSMD4/Rpn10, and ADRM1/Rpn13^2–5^. These substrates are then unfolded by translocating through the AAA-ATPase motor at the base of the 19S RP before entering the core particle for proteolysis^1,6,7^. Successful proteasomal degradation also involves PSMD14/Rpn11, an essential zinc metalloprotease on the 19S RP that deubiquitylates substrates to promote their efficient translocation^8–10^.

Some proteins can undergo proteasomal degradation without a requirement for ubiquitylation^11–15^. For example, the proteasomal adaptor midnolin directly captures specific nuclear proteins to promote their degradation^12,13^. It is unclear if and how other proteins engage proteasomes for ubiquitin-independent degradation. In this study, we report the cryo-EM structure of the human proteasome bound to thioredoxin-like protein 1 (TXNL1), a conserved thioreductase that interacts stably with proteasomes and shows genetic interactions with proteasome regulators in *S. pombe*^16–18^. Our structure reveals key binding interfaces between TXNL1 and proteasomal subunits required for the ubiquitin-independent degradation of TXNL1 upon cellular exposure to certain compounds that cause oxidative stress. Our findings identify the molecular interactions required for the stress-induced degradation of an abundant protein that may regulate proteasomal activity^18^.

We determined the structure of the TXNL1-bound proteasome to an overall resolution of 3.0-3.3 Å from cryo-EM datasets of affinity-purified midnolin-proteasome complexes (Fig. 1a, Extended Data Fig. 1,2, and Table 1). In addition to revealing structures of midnolin-bound proteasomes described elsewhere, 3D classifications revealed a subset of particles contributing to reconstructions of the proteasome or 19S RP with a well-resolved density that has not been previously reported (Fig. 1a and Extended Data Fig. 1, 2). We identified this protein as TXNL1 based on amino acid sequences assigned to the density by ModelAngelo^19^.

**Figure 1.**
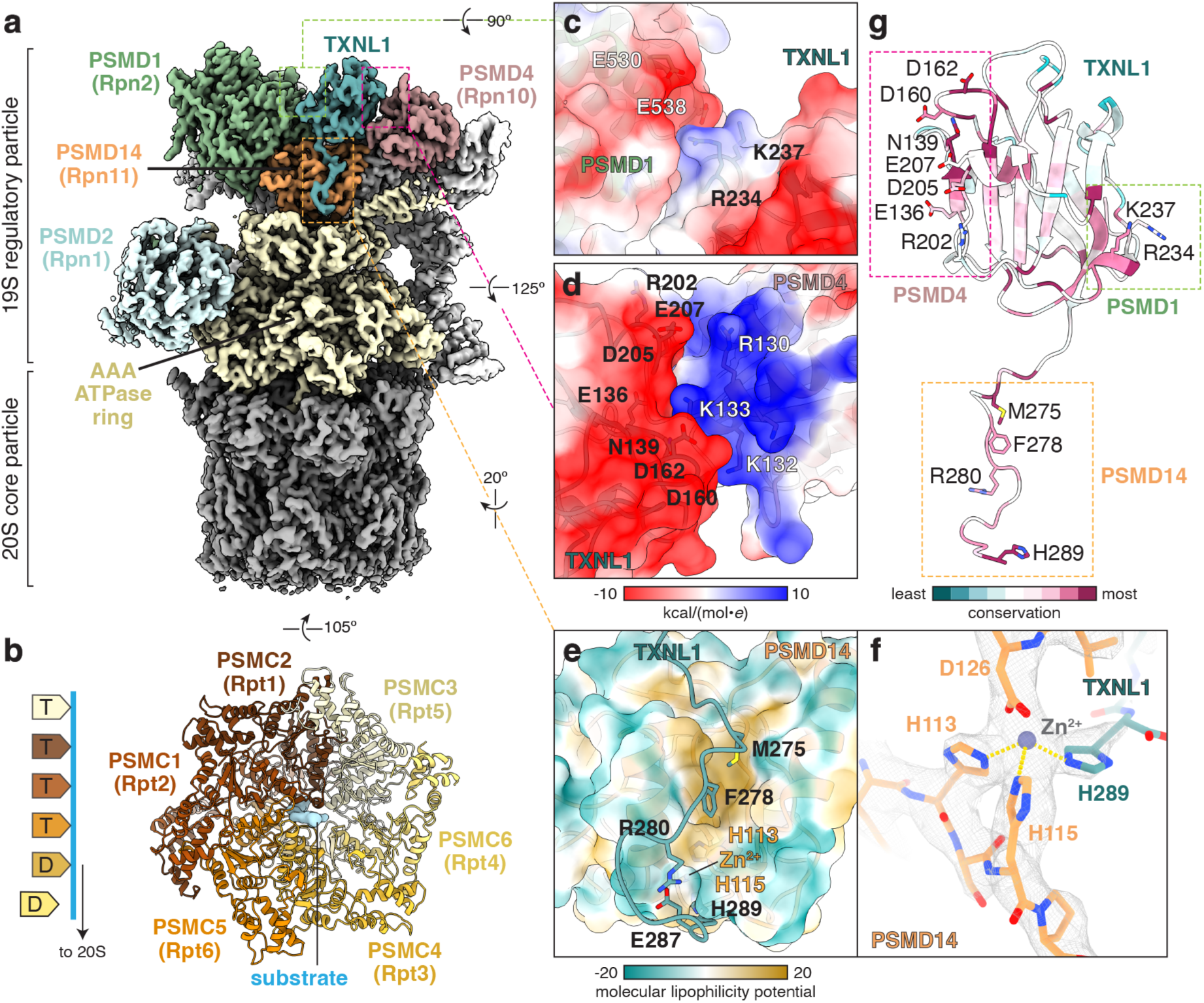
Structure of TXNL1 bound to the proteasome. **a**, Cryo-EM map of TXNL1 on the proteasome. **b,** Model of the AAA-ATPase ring with density corresponding to a translocating substrate (light blue), viewed from the 20S core particle (right), and scheme (left) depicting the configuration and nucleotide (T – ATP, D – ADP)-bound states of the ATPase subunits, colored as in the model, relative to the substrate. TXNL1 binding to **c,** PSMD1 and **d,** PSMD14 occurs through electrostatic interactions. **e,** The C-terminal tail of TXNL1 interacts with PSMD14 through hydrophobic interactions. **f,** H289 of TXNL1 coordinates the catalytic zinc ion associated with PSMD14. **g,** The proteasome-interacting domain of TXNL1 colored by conservation with residues at proteasomal interfaces labeled.

**Table 1.** Cryo-EM data collection, refinement, and validation statistics.

|  | TXNL1 +<br>proteasome<br>PDB 9BW4<br>EMD-44952 | TXNL1 +<br>proteasome (local<br>refinement)<br>EMD-44949 |
| --- | --- | --- |
| **Data collection and processing** |  |  |
| Magnification | 105,000 | 105,000 |
| Voltage (kV) | 300 | 300 |
| Electron exposure (e-/Å<sup>2</sup>) | 50 to 54 | 50 to 54 |
| Defocus range (μm) | -0.8 to -2.2 | -0.8 to -2.2 |
| Pixel size (Å) | 0.825 | 0.825 |
| Symmetry imposed | C1 | C1 |
| Initial particle images (no.) | 1,765,798 | 1,765,798 |
| Final particle images (no.) | 103,163 | 103,163 |
| Map resolution (Å) | 3.3 | 3.0 |
| FSC threshold | 0.143 | 0.143 |
| Map resolution range (Å) | 2.9 to 51.6 | 2.5 to 25.0 |
| **Refinement** |  |  |
| Initial model used (PDB code) | AlphaFold2,<br>ModelAngelo |  |
| Model resolution (Å) | 3.8 |  |
| FSC threshold | 0.5 |  |
| Model resolution range (Å) | 3.8 to 82.9 |  |
| Map sharpening *B* factor (Å<sup>2</sup>) | -55.6 |  |
| Model composition |  |  |
| Non-hydrogen atoms | 90,422 |  |
| Protein residues | 11,464 |  |
| Ligands | 1 Zn<sup>2+</sup><br>5 Mg<sup>2+</sup><br>4 ATP<br>2 ADP |  |
| *B* factors (Å<sup>2</sup>) |  |  |
| Protein | 108.57 |  |
| Ligand | 114.09 |  |
| R.m.s. deviations |  |  |
| Bond lengths (Å) | 0.005 |  |
| Bond angles (°) | 1.000 |  |
| Validation |  |  |
| MolProbity score | 2.26 |  |
| Clashscore | 15.28 |  |
| Poor rotamers (%) | 0.02 |  |
| Ramachandran plot |  |  |
| Favored (%) | 89.20 |  |
| Allowed (%) | 10.75 |  |
| Disallowed (%) | 0.04 |  |

TXNL1 is a 32 kDa protein with an N-terminal thioredoxin domain and a C-terminal proteasome interacting PITH (<u>p</u>roteasome-interacting <u>th</u>ioredoxin) domain^16–18^ (Extended Data Fig. 3a). Although the thioredoxin domain is not resolved in our map, we could model the entire PITH domain (residues 118-289) bound to PSMD1, a structural component of the 19S RP, the ubiquitin receptor PSMD4, and the deubiquitinase PSMD14. Below the TXNL1 binding site, the AAA-ATPase ring is in a state of active translocation, with PSMC3/Rpt5 positioned at the top of the spiral AAA-ATPase configuration, PSMC6/Rpt4 disengaged from the substrate, and nucleotide density associated with each subunit (Fig. 1b and Extended Data Fig. 3b). This configuration is similar to the human E_D2_ (RMSD = 3.0 Å) and yeast 4D (RMSD = 2.0 Å) states of proteasomes reported previously^6,7^ (Extended Data Fig. 3c).

Electrostatic interactions facilitate TXNL1 binding to PSMD1 and PSMD4 (Fig. 1c,d). Residues involved in these interactions include R234 of TXNL1, which interacts with an acidic patch on PSMD1 (Fig. 1c), and E136 and D162 of TXNL1, which interact with a basic interface on PSMD4 (Fig. 1d). The C-terminal unstructured tail of TXNL1 engages a hydrophobic groove on PSMD14 (Fig. 1e) and extends into the PSMD14 active site, where H289, the C-terminal residue of TXNL1, coordinates zinc along with H113 and H115 of PSMD14 (Fig. 1f,g). TXNL1 binding to PSMD14 would clash with ubiquitin, and when bound to TXNL1, the ubiquitin-interacting Insert-1 region of PSMD14 does not assume the β-hairpin conformation used to position ubiquitin for cleavage^6,7,20^ (Extended Data Fig. 3d). The amino acids on TXNL1 that interact with proteasomal subunits, particularly H289, are highly conserved (Fig. 1g).

TXNL1 is abundant (over 1 µM) and nearly equimolar with 19S RP subunits in HEK-293T cells^21^, and native size fractionations of HEK-293T lysates revealed that approximately 50% of endogenous TXNL1 co-migrated with proteasomes (Extended Data Fig. 4a). In addition, immunofluorescence showed TXNL1 enriched in the nucleus (Extended Data Fig. 4b,c) despite lacking a conventional nuclear localization sequence. Mutations at proteasome-binding interfaces or that disrupt thioreductase activity do not visibly change the localization of TXNL1 (Extended Data Fig. 4c). Considered together with the nuclear localization of midnolin^13^, the abundance and high proteasomal occupancy of TXNL1 may explain the presence of TXNL1-bound proteasomes in our datasets.

To validate the interaction interfaces observed in our structure, we performed co-immunoprecipitations from *TXNL1* knockout (KO) HEK-293T cells complemented with epitope-tagged TXNL1 variants (Fig. 2a). Wild type (WT) TXNL1 associated with proteasomes, but the interaction was severely impaired by mutations to disrupt the interface with PSMD1 (R234D; lane 10), PSMD4 (E136R and D162R; lanes 3-6), or PSMD14 (lanes 11-14). With the exception of N139A TXNL1 (lane 4), we observed an inverse correlation between TXNL1 abundance and its ability to bind proteasomes, consistent with the previous observation that TXNL1 levels are increased upon proteasome inhibition^16^. These interfaces match Alphafold3 predictions of 19S RP subunits with TXNL1 and PITHD1, which has a PITH domain but no thioreductase activity (Extended Data Fig. 5a,b). PITHD1 also has an extended C terminus that appeared to enter the central pore of the AAA-ATPase ring in the predictions. Although PITHD1 isolated proteasomes less efficiently than TXNL1, immunoprecipitations confirmed that mutations in the conserved interaction interfaces abolished proteasome binding (Extended Data Fig. 5c).

**Figure 2.**
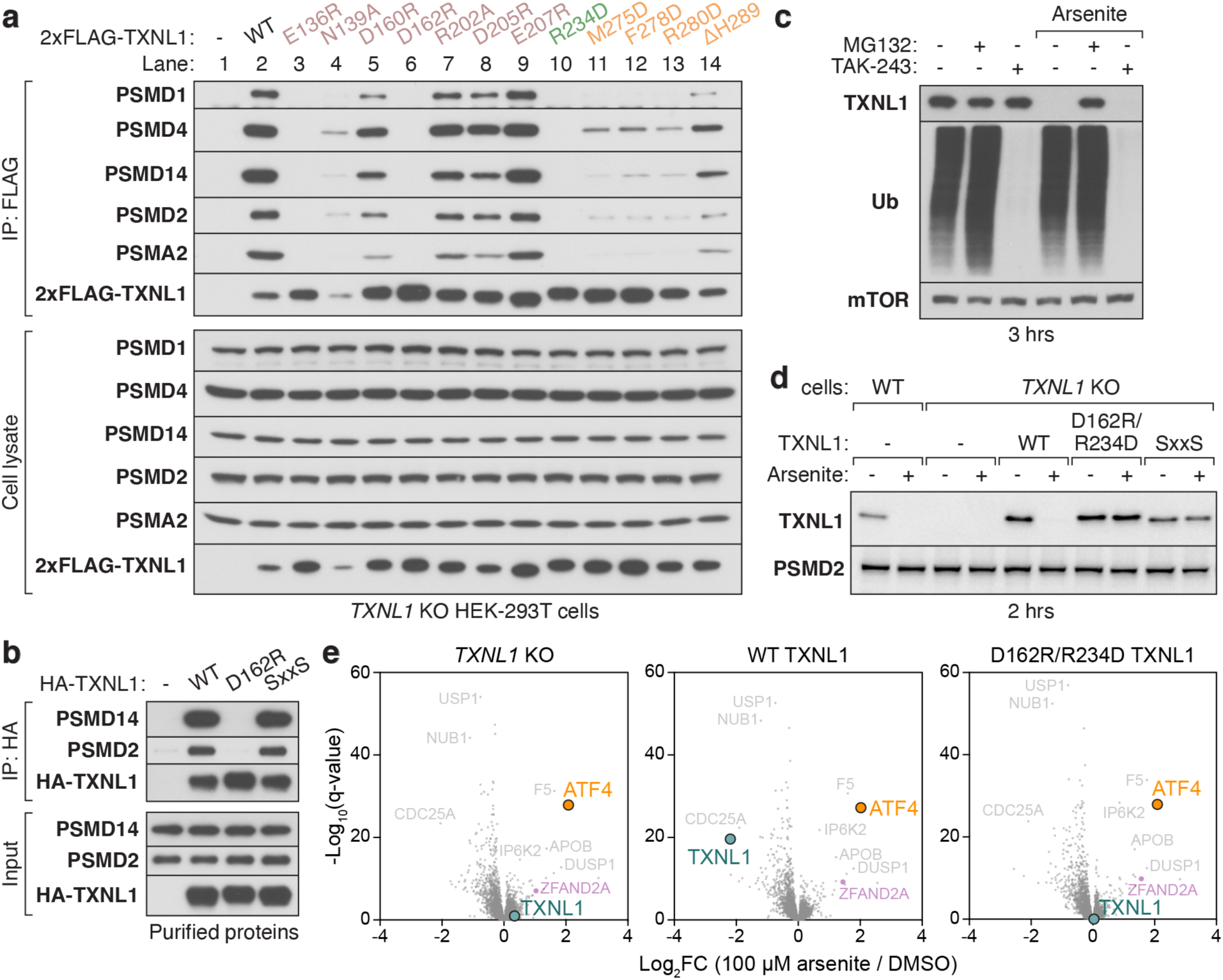
Proteasome binding is required for the stress-induced degradation of TXNL1. **a**, Immunoblotting of anti-FLAG immunoprecipitates (IP, top) and input lysates (bottom) of *TXNL1* knockout (KO) HEK-293T cells transfected with 2xFLAG-tagged TXNL1 variants. WT, wild type. **b,** Immunoblotting of purified HA-tagged TXNL1 variants immunoprecipitated (IP) in the presence of purified proteasomes. SxxS, thioreductase mutant. **c,** TXNL1 degradation upon arsenite treatment is proteasome-dependent but ubiquitin-independent. Immunoblotting of HEK-293T cells treated without or with 100 µM arsenite, 10 µM MG132, a proteasome inhibitor, or 1 µM TAK-243, an inhibitor of the E1 ubiquitin activating enzyme. Ub, ubiquitin. **d,** TXNL1 degradation upon arsenite exposure requires proteasome binding and thioreductase activity. Immunoblotting from WT or *TXNL1* KO HEK-293T cells stably complemented without or with WT and mutant versions of TXNL1 by lentiviral transduction. **e,** Volcano plots of showing the fold-change (Log_2_FC) of protein levels upon treatment with arsenite for 3 hrs of *TXNL1* KO HEK-293T cells complemented without or with WT or D162R/R234D TXNL1.

Next, we validated TXNL1-proteasome interactions with purified components. Immunoprecipitations of purified HA-TXNL1 variants incubated with proteasomes from *TXNL1* KO HEK-293T cells confirmed that WT TXNL1 and a thioredoxin domain mutant (SxxS) isolated proteasomes while D162R TXNL1, a proteasome binding-deficient mutant, did not (Fig. 2b and Extended Data Fig. 6a,b). TXNL1 binding to proteasomes was also impaired by pre-incubating proteasomes with 1,10-phenanthroline, a zinc chelator that inhibits PSMD14 deubiquitylation activity^8^, but not 1,7-phenanthroline, an inactive isomer (Extended Data Fig. 6c). This finding is consistent with our observation that H289 of TXNL1 coordinates the catalytic zinc of PSMD14 (Fig. 1f) and is important for proteasome engagement (Fig. 2a). However, unlike 1,10-phenanthroline, an excess of TXNL1 pre-incubated with purified proteasomes did not inhibit the *in vitro* deubiquitylation or degradation of a poly-ubiquitylated protein in an endpoint assay^22^ (Extended Data Fig. 6d).

It has been reported that exposing cells to arsenite, a chemotherapeutic used to treat acute promyelocytic leukemia that causes oxidative stress, reduces TXNL1 protein levels^17,23^. We confirmed that TXNL1 is rapidly destabilized in cells treated with arsenite but not hydrogen peroxide (Extended Data Fig. 7a,b). The proteasome inhibitor MG132 abrogated the arsenite-triggered degradation of TXNL1, but TAK-243, an inhibitor of the E1 ubiquitin-activating enzyme^24^, did not (Fig. 2c). Complementing *TXNL1* KO cells with TXNL1 variants showed efficient arsenite-induced degradation of WT TXNL1, but not of D162R/R234D TXNL1 deficient in proteasome binding or the SxxS thioredoxin mutant (Fig. 2d and Extended Data Fig. 7c). We observed similar results using auranofin, a gold-containing oxidizing agent^25–27^, in place of arsenite (Extended Data Fig. 7d,e). These observations indicate that metal- or metalloid-containing oxidative agents trigger a cellular response leading to the ubiquitin-independent degradation of TXNL1 that requires both the proteasome binding activity and catalytic cysteines of TXNL1.

To further investigate this response and the general role of TXNL1, we used multiplexed proteomics to analyze *TXNL1* KO cells complemented without or with WT or D162R/R234D TXNL1 (Fig. 2e, Extended Data Fig. 7f, and Supplementary Table 1). We also performed RNA sequencing of the same cell lines as well as *TXNL1* KO cells complemented with SxxS TXNL1 (Extended Data Fig. 8 and Supplementary Table 2). These datasets revealed that re-expressing any TXNL1 variant in KO cells minimally impacted the proteome and transcriptome (Extended Data Fig. 7f and 8a). Supporting our immunoblotting results, arsenite treatment selectively destabilized WT but not D162R/R234D TXNL1 protein levels and induced a dramatic transcriptional response and the posttranscriptional upregulation of the stress-induced transcription factor ATF4 in all cell lines (Fig. 2e and Extended Data Fig. 8b). However, we did not observe specific differences in the arsenite-dependent upregulation or downregulation of any protein or transcript besides TXNL1 across the cell lines. Thus, elucidation of the purpose and impact of TXNL1 degradation in cellular physiology remains to be determined.

Altogether, our study reveals specific interactions required for proteasome binding and the stress-induced degradation of TXNL1. Based on the high abundance of TXNL1, its thioreductase activity, and its position above the entry to the AAA-ATPase motor of actively translocating proteasomes, it is tempting to speculate that TXNL1 functions to reduce oxidized substrate proteins to facilitate their degradation. In this model, TXNL1 may cooperate with deubiquitylation by PSMD14 to promote efficient substrate translocation, which may involve TXNL1 coming on and off the proteasome when substrate-conjugated ubiquitins engage PSMD14 for removal. Because putative substrates that require TXNL1 for degradation are probably heterogeneous and constitute only a small fraction of proteins in a typical cell, like many proteins regulated by quality control pathways, identifying them may be challenging. Future insights, facilitated by the molecular interactions reported here, will require discovering specific substrates, cell types, or cellular conditions that are particularly reliant on TXNL1.

## Acknowledgments

Cryo-EM screening and data collection were performed at the Harvard Center for Cryo-Electron Microscopy (HC2EM). Data processing was supported by SBGrid. We thank H. Li and T. Rapoport for providing the poly-ubiquitylated protein for degradation assays, and members of the Elledge and Shao labs for useful discussions. This work was supported by a Robin Reed Memorial Fellowship (JG), a National Science Foundation Graduate Research Fellowship (CN), an American Heart Association predoctoral fellowship (MCJY), a Takeda Foundation Fellowship (HC), the National Mah Jongg League Fellow of the Damon Runyon Cancer Research Foundation (DRG-2469-22; XG), NIH R01GM132129 (JAP), a Packard fellowship (SS), NIH R01AG073277 (SS), and NIH AG11085 (SJE). SJE is a member of the Ludwig Center at Harvard. SS and SJE are Investigators with the Howard Hughes Medical Institute.

## Competing Interests

SJE is a founder of and hold equity in TScan Therapeutics, MAZE Therapeutics, and Mirimus and serves on the scientific advisory board of TSCAN Therapeutics, Infinity Bio, and MAZE Therapeutics. All other authors declare no competing interests.

## Author Contributions

JG and MCJY prepared cryo-EM samples and collected and processed cryo-EM data. CN generated cell lines and performed most cellular and biochemical experiments with help from HC, XG, and MJR. JG, MCJY, and SS modeled and interpreted the structure. HC prepared samples for proteomics analysis. CN and XG purified proteasomal complexes used for cryo-EM. CN and ZM performed RNA-Seq analysis. JAP oversaw proteomics analysis. CN, JG, SJE, and SS wrote the paper with input from all authors. SJE and SS supervised the project.

## Extended Data Figures

**Extended Data Figure 1.**
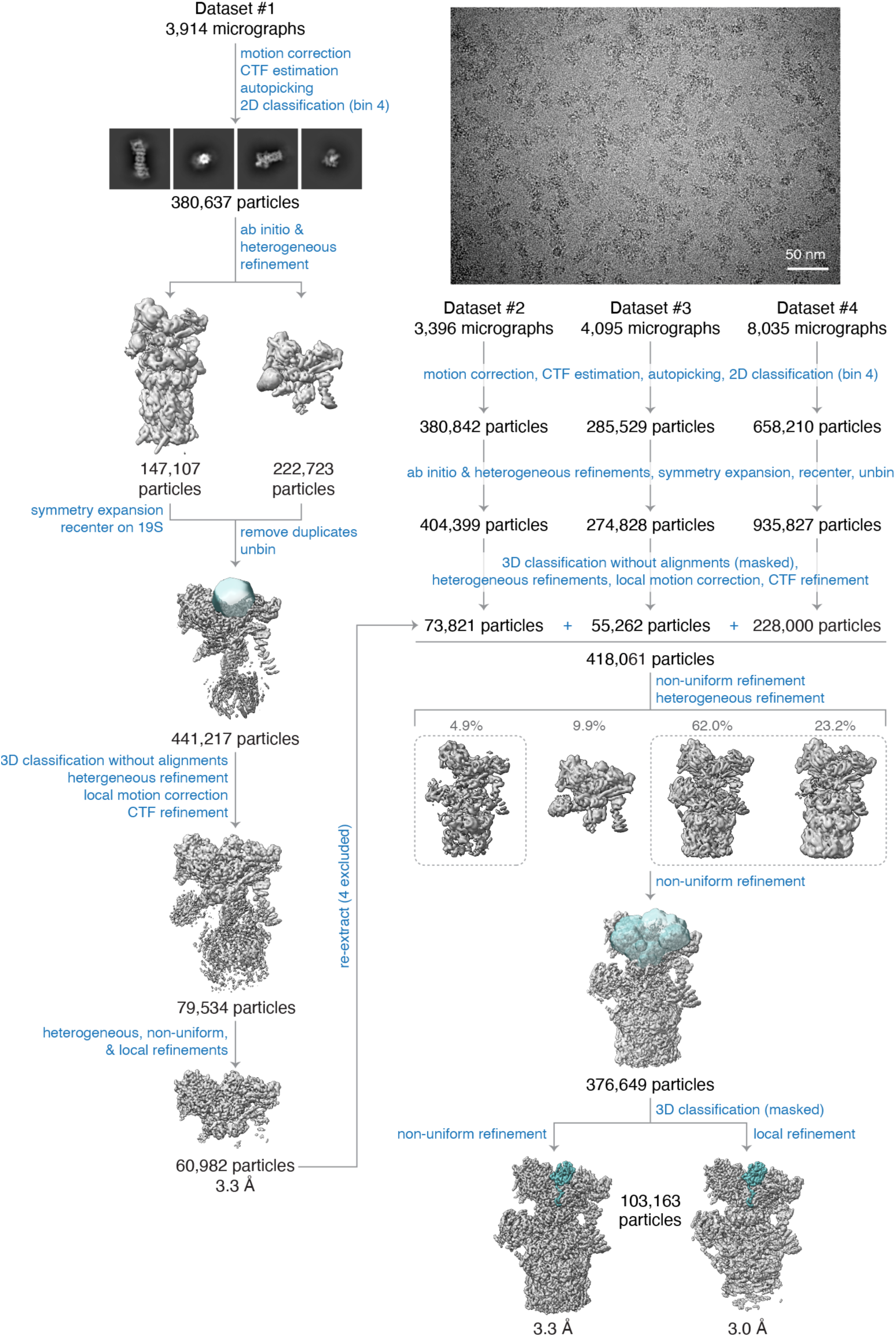
Cryo-EM data processing. Summary of cryo-EM data processing and classification strategy showing a representative micrograph, 2D class images, and 3D reconstructions.

**Extended Data Figure 2.**
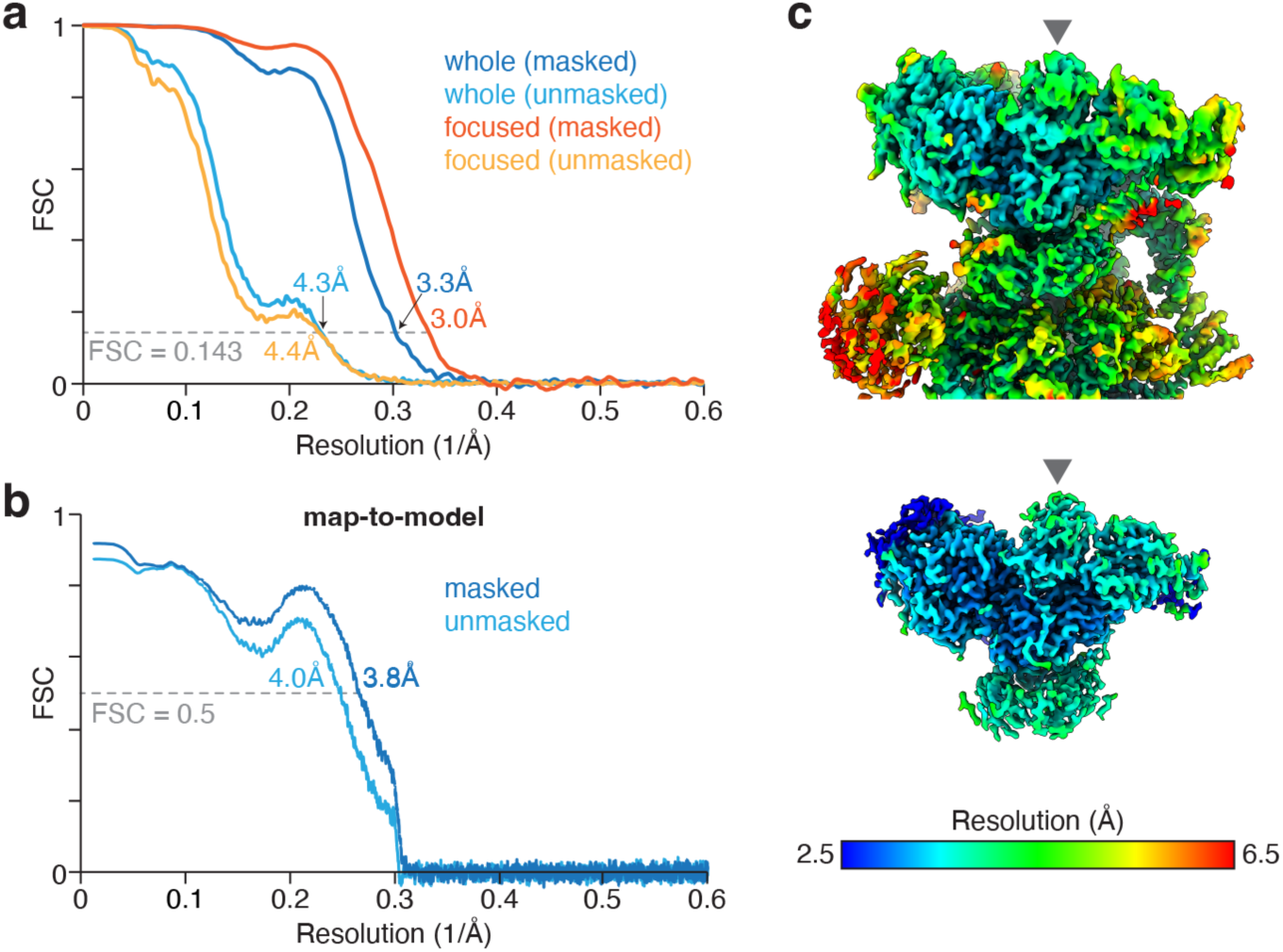
Cryo-EM map and model quality. **a**, Fourier shell correlation (FSC) vs. resolution (1/Å) curves for the indicated cryo-EM maps. **b,** The whole (top) and focused (bottom) cryo-EM map colored by local resolution. Arrow denotes TXNL1. **c,** Model vs. map FSC curves.

**Extended Data Figure 3.**
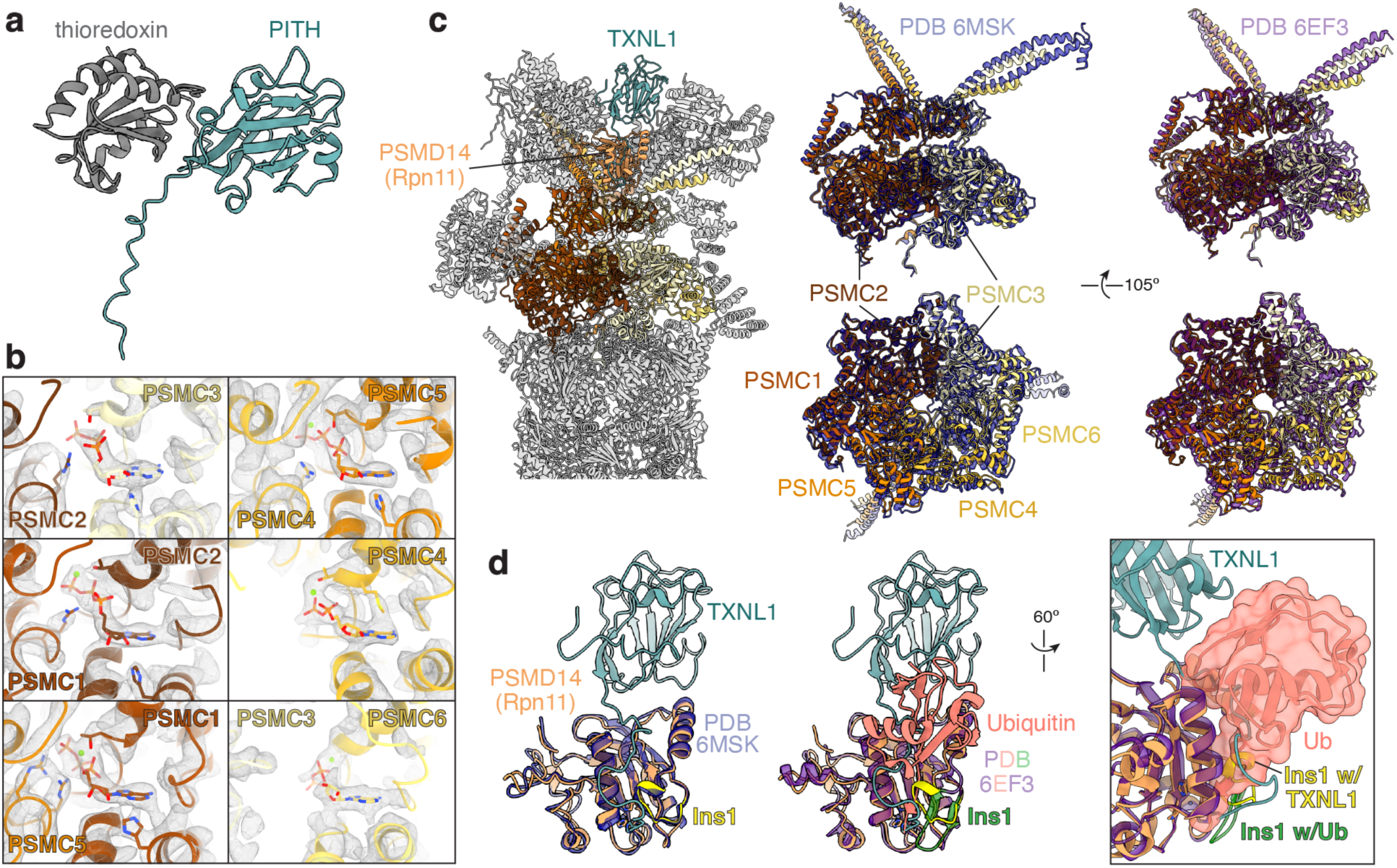
Characteristics of the TXNL1-bound proteasome. **a**, AlphaFold prediction of TXNL1 showing the N-terminal thioredoxin domain and C-terminal proteasome-interacting thioredoxin (PITH) domain. **b,** Map density and model of the nucleotide binding sites in the AAA-ATPase ring. **c,** Structure of the TXNL1-bound proteasome with TXNL1, PSMD14, and AAA-ATPase subunits colored as in Fig. 1b (left) and superposition of the AAA-ATPase ring with that in the E_D2_ state of the human proteasome (PDB 6MSK; transparent dark blue; middle) or the 4D state of the yeast proteasome (PDB 6EF3; transparent dark purple, right). **d,** TXNL1 engagement with PSMD14 is incompatible with ubiquitin binding. Superposition of PSMD14 bound to TXNL1 colored as in **c** with PSMD14 (PDB 6MSK, transparent dark blue, left) or ubiquitin (salmon; Ub)-bound Rpn11 (PDB 6EF3, transparent purple, right). The Insert-1 (Ins1) region of PSMD14/Rpn11 that assumes a β-hairpin when bound to ubiquitin is colored yellow when bound to TXNL1 and green when bound to ubiquitin. Note: ubiquitin binding to PSMD14 would clash with the TXNL1 C-terminal tail and the TXNL1-bound conformation of Insert-1.

**Extended Data Figure 4.**
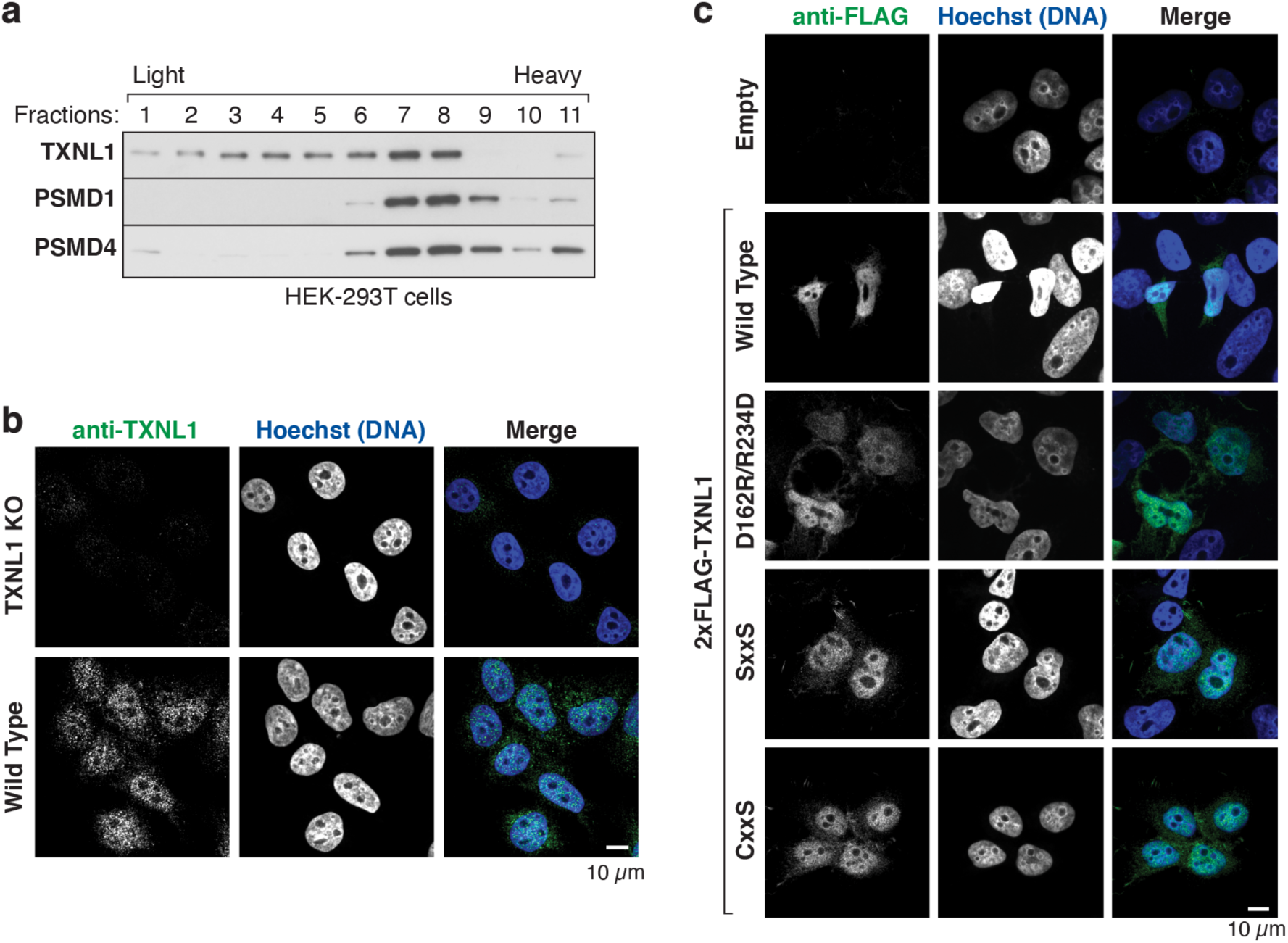
TXNL1 localization and proteasomal occupancy. **a**, TXNL1 co-migrates with proteasomal components in size fractionations of HEK-293T cell lysates by sucrose gradient ultracentrifugation, assayed by SDS-PAGE and immunoblotting. **b,** Endogenous TXNL1 is localized in the nucleus. anti-TXNL1 immunofluorescence of wild type and *TXNL1* knockout (KO) HEK-293T cells. **c,** anti-FLAG immunofluorescence of *TXNL1* KO HEK-293T cells with transient transfection of the indicated 2xFLAG-tagged TXNL1 variants. The D162R/R234D mutant disrupts proteasome binding and the SxxS and CxxS mutants disrupt thioreductase activity.

**Extended Data Figure 5.**
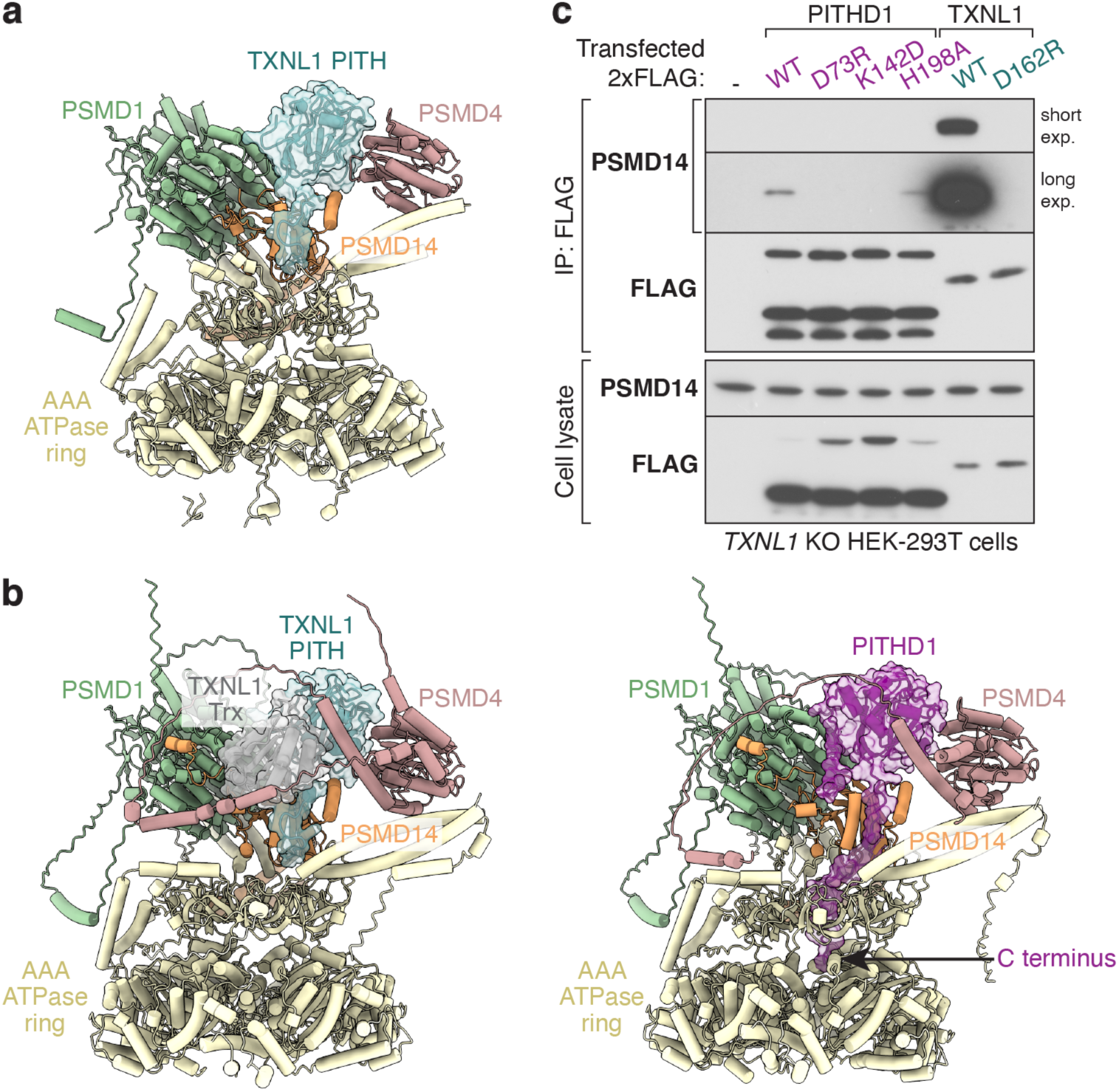
Common PITH domain interaction interfaces. **a**, Cryo-EM structure showing TXNL1 PITH domain interactions with the proteasome. **b,** Alphafold3 models of TXNL1 (left) or PITHD1 (right) with 19S regulatory particle subunits: PSMD1, PSMD4, PSMF14, and the AAA-ATPase subunits show shared interaction interfaces. Note: PITHD1 has an extended C-terminal tail past the histidine predicted to coordinate the zinc on PSMD14 that appears to extend into the central pore of the AAA-ATPase ring. **c,** Immunoblotting of anti-FLAG immunoprecipitates (IP, top) and input lysates (bottom) of *TXNL1* knockout (KO) HEK-293T cells transfected with 2xFLAG-tagged PITHD1 and TXNL1 variants. WT, wild type. D73 of PITHD1 is equivalent to D162 of TXNL1, K142 of PITHD1 is equivalent to R234 of TXNL1, and H198 of PITHD1 is equivalent to H289 of TXNL1, but is followed by a 13-residue extension.

**Extended Data Figure 6.**
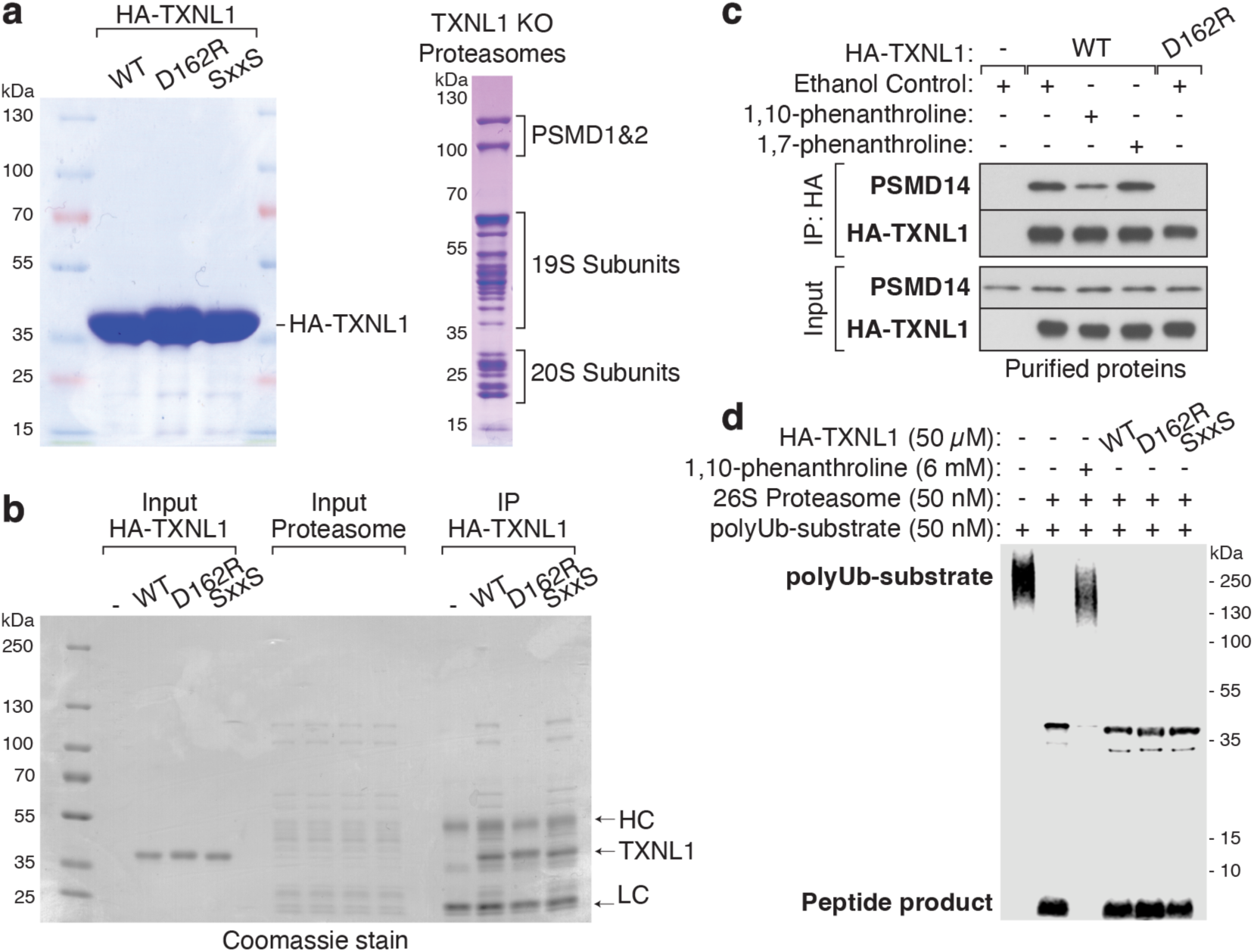
*In vitro* assays of TXNL1. **a**, SDS-PAGE and Coomassie staining of purified HA-tagged TXNL1 variants (left) and TXNL1-deficient human proteasomes (right). **b,** Coomassie stain of samples from Fig. 2b. IP, immunoprecipitation. **c,** Purified HA-TXNL1 variants were immunoprecipitated in the presence of purified proteasomes pre-incubated with 6 mM of the indicated compound for 30 min at room temperature before immunoblotting. **d,** Proteasomes were pre-incubated with TXNL1 or the PSMD14/Rpn11 inhibitor 1,10-phenanthroline for 30 min at room temperature before the addition of a polyubiquitylated (polyUb) sfGFP protein substrate for 10 min at 37°C. Note: 1,10-phenanthroline inhibits deubiquitylation and degradation of the polyUb-substrate (lane 3), but an excess of TXNL1 does not (lane 4).

**Extended Data Figure 7.**
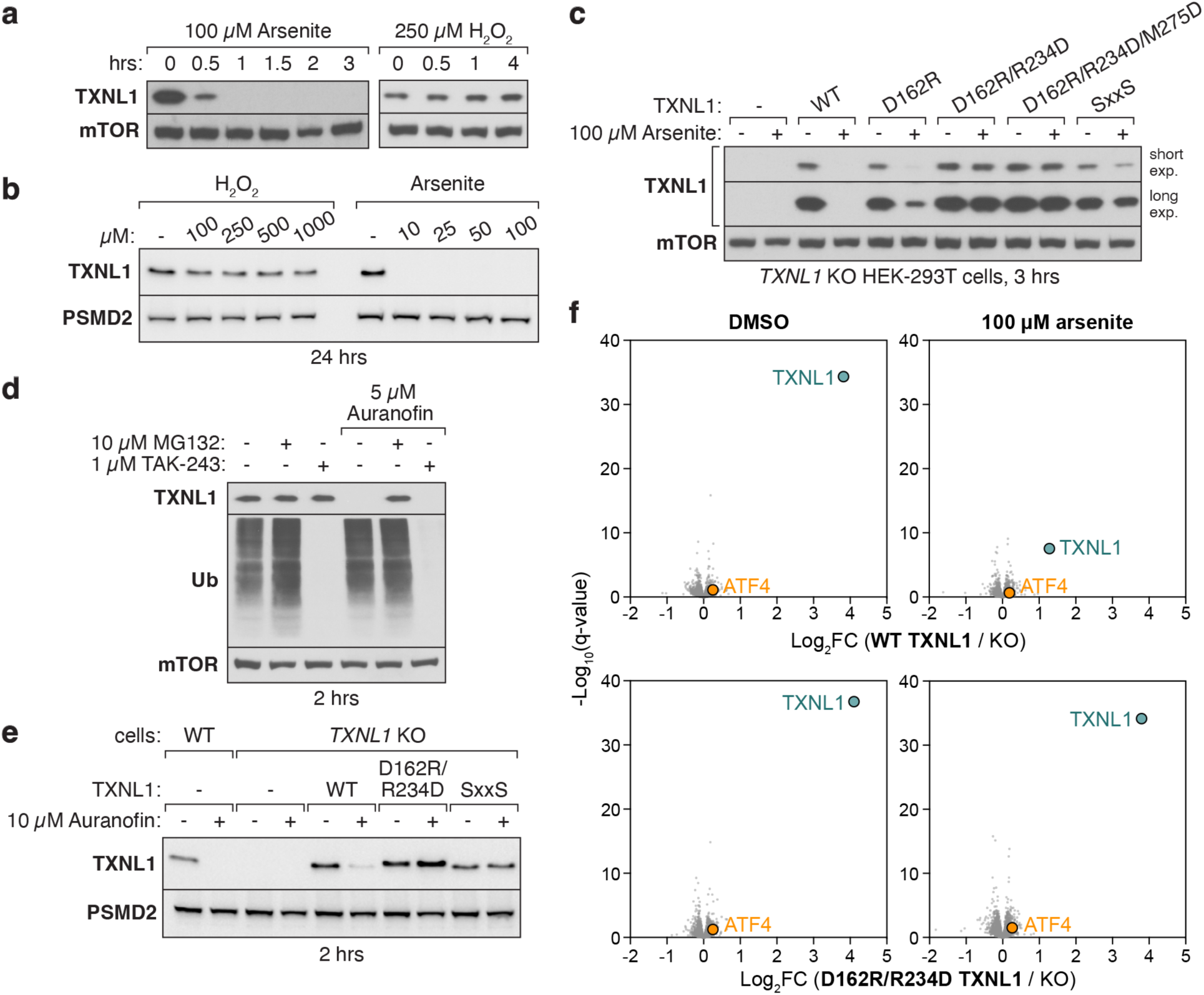
Requirements for stress-induced TXNL1 degradation. **a,b**, TXNL1 is rapidly degraded upon cellular exposure to arsenite but not hydrogen peroxide. Immunoblotting of HEK-293T cells treated with the indicated concentration of oxidative agents for the time periods indicated. **c,** Immunoblotting from *TXNL1* knockout (KO) HEK-293T cells that were stably complemented with wild type (WT) and mutant versions of TXNL1 by lentiviral transduction. D162, R234, and M275 reside at the interaction interfaces with PSMD1, PSMD4, and PSMD14, respectively. SxxS mutates the catalytic cysteines in the thioredoxin domain of TXNL1. Note: D162R TXNL1 does not co-purify proteasomes (Fig. 2b) but only weakly impairs arsenite-induced degradation, indicating that it still has some affinity for proteasomes, likely through the interface with PSMD1 that is disrupted by the R234D mutation. **d,e,** TXNL1 degradation upon auranofin treatment requires proteasome binding and thioredoxin activity. Assays as in Fig. 2c**,d** but with auranofin treatment as indicated in place of arsenite treatment. **f,** Selective degradation of TXNL1 upon arsenite exposure requires proteasome binding. Volcano plots of multiplexed proteomics data showing the fold-change (Log2FC) of protein levels in *TXNL1* KO HEK-293T cells complemented with the indicated TXNL1 variant compared to KO cells without (left) and with (right) arsenite treatment. Note: arsenite treatment selectively lowers the level of WT but not D162R/R234D TXNL1.

**Extended Data Figure 8.**
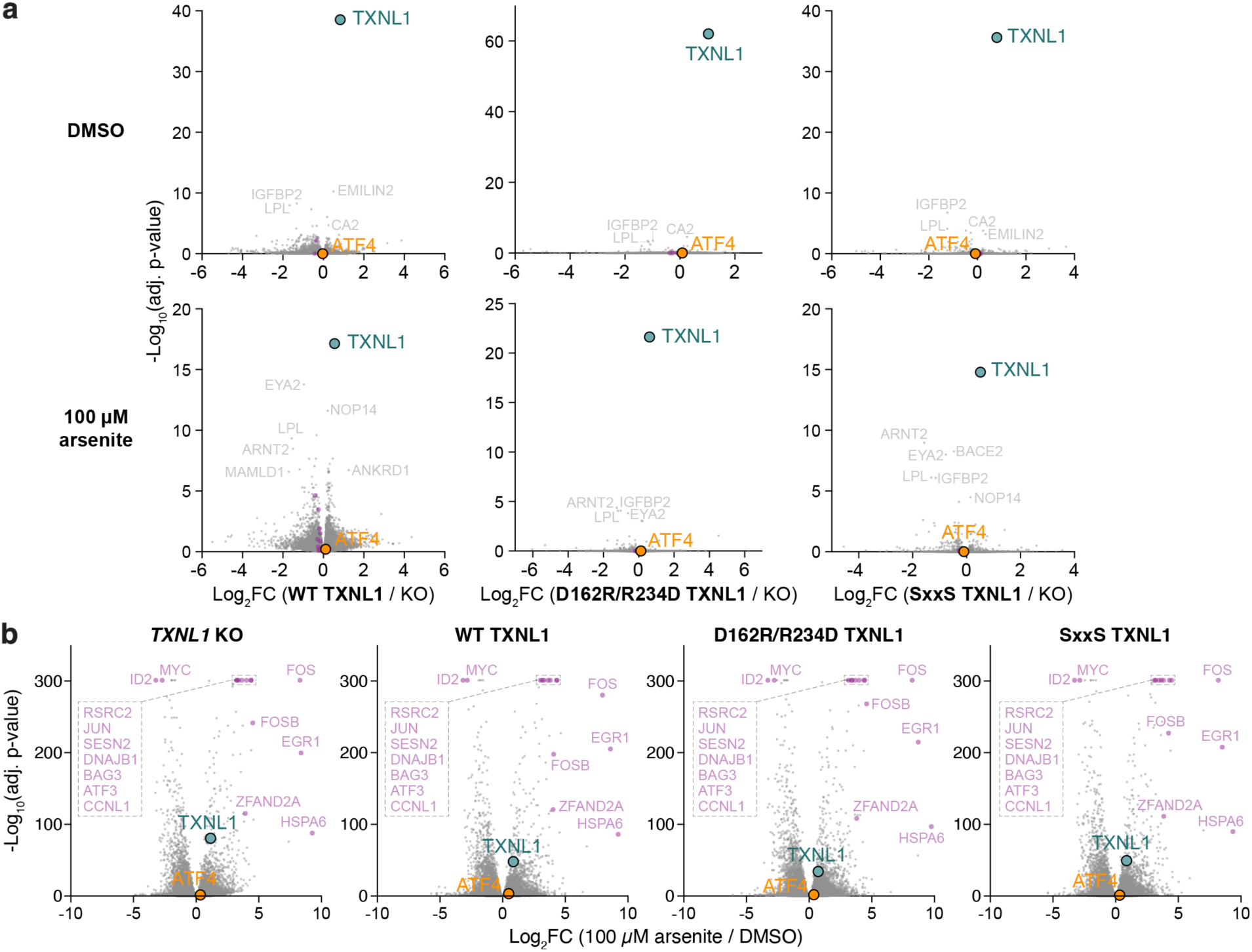
Transcriptional changes upon cellular arsenite exposure. **a,** Volcano plots of RNA sequencing data showing the fold-change (Log_2_FC) of transcript levels in *TXNL1* knockout (KO) HEK-293T cells complemented with wild type (WT) TXNL1, D162R/R234D TXNL1 deficient in proteasome binding, or SxxS TXNL1 deficient in thioreductase activity compared to KO cells without (top) or with (bottom) arsenite treatment. Note: TXNL1 is the main transcript that is significantly changes in both conditions. **b,** Volcano plots of RNA sequencing data showing the fold-change (Log_2_FC) of transcript levels in *TXNL1* KO cells without (left) or with complementation with WT TXNL1 or the indicated TXNL1 mutants upon arsenite treatment. Note: changes in stress-responsive transcript (purple) levels are not impacted by TXNL1 expression.

## Materials and Methods

### Cryo-EM sample preparation, data collection, image processing, and model building

Plasmid DNA encoding epitope-tagged midnolin variants was transiently transfected into human embryonic kidney (HEK)-293T cells for immunoprecipitation. The affinity purifications were partitioned by size-exclusion chromatography and the fractions corresponding to proteasomes were collected. 3.5 µl of 4 mg/mL proteasomal complexes was applied to glow-discharged R0.6/1 UltrAuFoil 300 mesh grids (Quantifoil) and frozen in liquid ethane using a Vitrobot Mark IV (Thermo Fisher Scientific) set at 4°C and 100% humidity with a 10 sec wait time, 3 sec blot time, and +8 blot force.

Datasets were collected using a Titan Krios (Thermo Fisher Scientific) operating at 300 kV and equipped with a BioQuantum K3 imaging filter with a 20-eV slit width and a K3 summit direct electron detector (Gatan) in counting mode at a nominal magnification of 105,000× corresponding to a calibrated pixel size of 0.825 Å. Semi-automated data collection was performed with SerialEM v4.0.5 for four datasets with the following parameters. Dataset #1: 2.597-second exposures were fractionated into 50 frames, resulting in a total exposure of 52.98 e-/Å^2^. The defocus targets were −1.0 to −2.2 µm. Dataset #2: 2.652-second exposures were fractionated into 48 frames, resulting in a total exposure of 50.54 e-/Å^2^. The defocus targets were −1.0 to −2.2 µm. Dataset #3: 2.65-second exposures were fractionated into 50 frames, resulting in a total exposure of 53 e-/Å^2^. The defocus targets were −1.0 to −2.2 µm. Dataset #4: 3.02-second exposures were fractionated into 54 frames, resulting in a total exposure of 53.79 e-/Å2. The defocus targets were −0.8 to −1.8 µm.

Data processing was performed in cryoSPARC v4.3.1. After patch-based motion correction and CTF estimation, micrographs with severe contamination or poor CTF fits were removed. 3,914 (Dataset #1), 3,396 (Dataset #2), 4,095 (Dataset #3) or 8,035 (Dataset #4) micrographs were subjected to automated particle picking using templates generated from blob-based picking. The particles were extracted with a box size of 800 and downsampled to a box size of 200. After 2D classification, 380,637 (Dataset #1), 380,842 (Dataset #2), 285,529 (Dataset #3) and 658,210 (Dataset #4) particles were selected for heterogeneous refinement using multiple reference volumes generated by ab initio reconstruction. Particles in the best classes were subjected to homogeneous refinement with C2 symmetry, followed by symmetry expansion. Afterwards, all particles were re-centered on the 19S particle, extracted with a box size of 440 and downsampled to a box size of 220. Good 19S particles after 2D classification were then subjected to heterogeneous refinements and 3D classification without alignment using a mask over the TXNL1 binding site based on initial observation of density in this region from heterogeneous refinements of Dataset #1 (Extended Data Fig. 1).

418,061 particles from the four datasets in classes with density corresponding to the TXNL1 were unbinned, local motion corrected, combined, and subjected to non-uniform and an additional round of heterogeneous refinement. Classes corresponding to intact proteasomes were selected and subjected to non-uniform refinement and 3D classification with a mask focused on TXNL1. 103,163 particles with clear TXNL1 density were used for final non-uniform and local refinements using the same mask. TXNL1 was identified by BLASTing the sequence assigned to this density by ModelAngelo^19^. An initial model for TXNL1-proteasome complex was generated primarily by rigid body fitting individual Alphafold models^28^ of each proteasomal subunit and of TXNL1, in addition to cross-referencing the output from ModelAngelo. The model was manually adjusted in Coot^29^ v0.9.8 in between iterative rounds of real space refinement in Phenix^30^ v1.19.2. Figures were made using ChimeraX^31^ v1.5 – v1.7.

### Expression plasmids

The human TXNL1 cDNA oligonucleotide, flanked with attB1/2 sites, was synthesized by Integrated DNA Technologies (IDT) for Gateway cloning into pDONR221 via a BP reaction (Thermo Fisher Scientific, 11789020). Mutations to the TXNL1 entry clone were introduced using the Q5 Site-Directed Mutagenesis kit (NEB, E0554S). The PCR primers were designed using the NEBaseChanger program and synthesized by IDT. Wild type and mutant versions of TXNL1 were subcloned into a pHAGE CMV 2xFLAG-N destination vector for N-terminal tagging or a pHAGE CMV destination vector conferring puromycin resistance^13^ via LR reactions (Thermo Fisher Scientific, 11791100).

HA-TXNL1 (WT, D162R, SxxS) were codon-optimized for expression in bacteria and synthesized by IDT to contain overhangs for Gibson assembly (NEB, E2611S). A pET 6xHis-3C bacterial expression vector was obtained from the DFCI Crystallography core facility and was digested with BamHI and NotI before Gibson assembly of the synthetic oligonucleotides.

The following proteins were expressed in bacteria:

HA-TXNL1 (WT):

MQLSHHHHHHSSGLEVLFQGPGMYPYDVPDYAGGGSVGVKPVGSDPDFQPELSGAGSRLAVVKFTMRGCGPCLRIAPAFSSMSNKYPQAVFLEVDVHQCQGTAATNNISATPTFLFFRNKVRIDQYQGADAVGLEEKIKQHLENDPGSNEDTDIPKGYMDLMPFINKAGCECLNESDEHGFDNCLRKDTTFLESDCDEQLLITVAFNQPVKLYSMKFQGPDNGQGPKYVKIFINLPRSMDFEEAERSEPTQALELTEDDIKEDGIVPLRYVKFQNVNSVTIFVQSNQGEEETTRISYFTFIGTPVQATNMNDFKRVVGKKGESH

HA-TXNL1 (D162R):

MQLSHHHHHHSSGLEVLFQGPGMYPYDVPDYAGGGSVGVKPVGSDPDFQPELSGAGSRLAVVKFTMRGCGPCLRIAPAFSSMSNKYPQAVFLEVDVHQCQGTAATNNISATPTFLFFRNKVRIDQYQGADAVGLEEKIKQHLENDPGSNEDTDIPKGYMDLMPFINKAGCECLNESDEHGFDNCLRKDTTFLESDCREQLLITVAFNQPVKLYSMKFQGPDNGQGPKYVKIFINLPRSMDFEEAERSEPTQALELTEDDIKEDGIVPLRYVKFQNVNSVTIFVQSNQGEEETTRISYFTFIGTPVQATNMNDFKRVVGKKGESH

HA-TXNL1 (SxxS):

MQLSHHHHHHSSGLEVLFQGPGMYPYDVPDYAGGGSVGVKPVGSDPDFQPELSGAGSRLAVVKFTMRGSGPSLRIAPAFSSMSNKYPQAVFLEVDVHQCQGTAATNNISATPTFLFFRNKVRIDQYQGADAVGLEEKIKQHLENDPGSNEDTDIPKGYMDLMPFINKAGCECLNESDEHGFDNCLRKDTTFLESDCDEQLLITVAFNQPVKLYSMKFQGPDNGQGPKYVKIFINLPRSMDFEEAERSEPTQALELTEDDIKEDGIVPLRYVKFQNVNSVTIFVQSNQGEEETTRISYFTFIGTPVQATNMNDFKRVVGKKGESH

### Cell culture and cell line generation

Human embryonic kidney (HEK)-293T cells (ATCC, CRL-3216, RRID: CVCL_0063) were incubated at 37°C and 5% CO_2_ in Dulbecco’s Modified Eagle’s Medium (DMEM) (Thermo Fisher Scientific, 11965118) supplemented with 100 units/mL of penicillin, 0.1 mg/mL of streptomycin (Thermo Fisher Scientific, 15070063), and 10% fetal bovine serum (Cytiva, SH30088.03). Cells were treated with 10 µM MG132 (Selleckchem, S2619), 1 µM TAK-243 (MedChemExpress, HY-100487), 100 µM arsenite (Sigma, S7400), 5 µM auranofin (MedChemExpress, HY-B1123), or 250 µM hydrogen peroxide (Sigma, 216763) from stock solution in dimethyl sulfoxide (DMSO) or water.

To knock out TXNL1, wild type HEK-293T cells were transfected using PolyJet (SignaGen, SL100688) with a lentiCRISPRv2 plasmid expressing a blue-fluorescent protein (*26*) and the guide RNA targeting TXNL1 (GAATGACCCTGGAAGCAATG). Five days post-transfection, the BFP-positive cells were single-cell sorted into 96-well plates and grown out for 2 weeks. Isogenic clones were expanded and screened for TXNL1 knockout.

To generate lentivirus, HEK-293T cells were cultured in 6-well plates to 90% confluency. Using PolyJet, the cells were transfected with 1 µg of total plasmid DNA encoding Tat, Rev, Gag-Pol, and VSV-G (mixed equimolar) and a lentiviral transfer vector. The media was replaced 1-day post-transfection. The lentiviral supernatant was collected 2 days post-transfection, passed through a 0.45 µm filter, and applied directly onto cells.

### Antibodies

The following primary antibodies were used for immunoblotting, all at a 1:1000 dilution: rabbit anti-PSMD14 (CST, 4197S, RRID: AB_11178935), rabbit anti-PSMD4 (CST, 12441S, RRID: AB_2797916), mouse anti-PSMD1 (Santa Cruz, sc-166038, RRID: AB_2172797), rabbit anti-PSMD2 (CST, 25430, RRID: AB_2798903), rabbit anti-PSMA2 (CST, 2455, RRID: AB_2171400), rabbit anti-FLAG (CST, 14793, RRID: AB_2572291), rabbit anti-HA (CST, 3724S, RRID: AB_1549585), rabbit anti-mTOR (CST, 2983, RRID: AB_2105622), rabbit anti-TXNL1 (Abcam, ab188328, RRID: AB_2687563), and rabbit anti-Ubiquitin (CST, 43124S, RRID: AB_2799235). Secondary antibodies used at a 1:2000 dilution were: anti-rabbit IgG, HRP-linked (CST, 7074, RRID: AB_2099233) or anti-mouse IgG, HRP-linked (CST, 7076S, RRID: AB_330924)

### Sucrose gradient ultracentrifugation

A stock solution containing 70% sucrose in water was first made. From this, stock solutions of 30%, 25%, 20%, 15%, and 10% sucrose were made in a final solvent concentration of 100 mM NaCl and 40 mM HEPES pH 7.4. Five 40 µL steps of 30%, 25%, 20%, 15%, and 10% sucrose in a matching buffer were added to an Eppendorf tube (200 µL total volume) and the gradient was left to sit for 30 min at 4°C. About 2 million HEK-293T cells were lysed in 50 µL of lysis buffer containing 0.5 % CHAPS, 100 mM NaCl, 40 mM HEPES pH 7.4, and 1x protease-phosphatase inhibitor cocktail. 20 µL of clarified lysate supernatant was loaded onto the gradient before ultracentrifugation at 55,000 rpm for 1 h using a TLS55 rotor. After centrifugation, 20 µL aliquots were diluted in Tris-Glycine sample buffer containing 10% 2-mercaptoethanol.

### Immunofluorescence

HEK-293T cells were seeded into 6-well plates containing Poly-D-Lysine coated coverslips (TED PELLA, Inc.) and grown to about 50% confluency. The cells were rinsed once with ice-cold PBS and were fixed with 4% paraformaldehyde (PFA) diluted in PBS for 15 min at room temperature. The cells were then rinsed thrice using ice-cold PBS and permeabilized with 0.05% Triton diluted in PBS for 5 min at room temperature. The cells were rinsed thrice using ice-cold PBS and incubated in immunofluorescence blocking buffer (LI-COR, 927-70001) for 45 min at room temperature. Coverslips were placed faceup on parafilm and incubated with rabbit anti-TXNL1 (Abcam, ab188328, RRID: AB_2687563) at 1:250 dilution in blocking buffer overnight. After incubation, the coverslips were rinsed thoroughly by dipping 50 times in PBS. Coverslips were again placed faceup on parafilm and incubated with blocking solution containing 1:500 dilution of anti-rabbit Alexa 488-conjugated secondary antibody (Thermo Fisher Scientific, A32731, RRID: AB_2633280) and 8 µM Hoechst 33342 dye (Thermo Fisher Scientific, H3570) for 1 h at room temperature. The coverslips were rinsed thoroughly by dipping 50 times in PBS and mounted on glass slides using ProLong Glass antifade mountant (Thermo Fisher Scientific, P36982).

Images were collected using a W1 Yokogawa Spinning disk Nikon Ti inverted confocal microscope with a 50 µm pinhole disk, a Plan Apo 60X/1.4 oil immersion objective, an Andor Zyla 4.2 Plus sCMOS monochrome camera, and excitation lasers 405 nm and 488 nm. Data was collected using the Nikon Elements Acquisition Software AR 5.02.

### Immunoprecipitations from cells

HEK-293T cells were seeded at 1 million cells in 10 cm culture dishes and transfected with 3 µg of DNA encoding 2xFLAG-TXNL1 variants using PolyJet 2 days post-seeding. The media was replaced with fresh media 1-day post-transfection and incubated overnight. The cells were rinsed once with ice-cold PBS and 700 µL of lysis buffer containing 0.5% CHAPS, 100 mM NaCl, 40 mM HEPES pH 7.4, 1x protease-phosphatase inhibitor cocktail (Thermo Fisher Scientific, 78441) was applied directly to the dish. The cells were collected into Eppendorf tubes by scraping and incubated for 15 min at 4°C with end-to-end rotation. The lysates were then centrifuged at 21,000 xg for 15 min at 4°C. While centrifuging, 15 µL/plate of anti-FLAG magnetic beads (Sigma, M8823, RRID: AB_2637089) were rinsed thrice with the same lysis buffer. After centrifuging, 30 µL of lysate was diluted into 200 µL of Tris-Glycine SDS sample buffer (Thermo Fisher Scientific, LC2676) supplemented with 10% 2-mercaptoethanol. The remaining supernatant was applied directly to the anti-FLAG beads and incubated end-to-end for 1 h at 4°C. After incubation, the beads were rinsed thrice with 700 µL of lysis buffer and resuspended in 50 µL of sample buffer. Proteins were eluted from the beads and denatured for 3 min at 95°C. An aliquot of IP sample was diluted 1:20 in sample buffer to blot for the abundant bait protein. To perform immunoblotting, 15 µL of lysate, diluted IP, or total IP was loaded into 15-well gels.

### Purification of recombinant HA-TXNL1

Rosetta (DE3) cells (Millipore, 71397-3) were transformed with pET plasmids encoding 6xHis-3C-HA-TXNL1 (WT, D162R, SxxS) and grown overnight at 37°C on plates containing kanamycin + chloramphenicol. Single colonies were then grown out in 50 mL of LB containing kanamycin + chloramphenicol overnight at 37°C with 200 rpm shaking. The following day, the LB-containing bacteria were transferred to 1 L of fresh LB solution containing kanamycin + chloramphenicol and grown at 37°C with 200 rpm shaking to an O.D. = 0.6. The bacteria were kept at 4°C for 10 min before supplementing the LB with 0.5 mM isopropyl ϕ3-d-1-thiogalactopyranoside IPTG (Sigma, I5502-10G) to induce TXNL1 expression. Cultures were incubated at 16°C for 16 h with 200 rpm shaking.

The following day, the liquid cultures were transferred to 1 L centrifuge bottles, and bacteria were pelleted by centrifuging at 4,000 xg for 15 min at 4°C. The pellets were resuspended in 35 mL of buffer containing 50 mM HEPES pH 7.4, 150 mM NaCl, and 10 mM imidazole (Sigma, 68268-500ML-F). The bacteria were again pelleted by centrifugation and the supernatant was discarded. The bacteria were then resuspended in 35 mL of lysis buffer containing 50 mM HEPES pH 7.4, 150 mM NaCl, 10 mM imidazole, 1 mM dithiothreitol DTT (Thermo Fisher Scientific, R0862), benzonase (Millipore, 71206-3), and protease inhibitor tablets (Sigma, 11873580001). A microfluidizer (700 bar, 4 cycles, 20 mL/min flow rate) was then used to lyse cells. The lysate was centrifuged at 21,000 xg for 20 min at 4°C. During the centrifugation, 4.8 mL of Ni-NTA agarose (Qiagen, 30210) was rinsed thrice using lysis buffer. Post-centrifugation, the supernatant was incubated with the washed Ni-NTA resin in 50 mL conical tubes for 30 min at 4°C with end-to-end rotation. The solution was then passed through a column (BioRad, 7321010) to remove the liquid by gravity. The conical tubes and column were rinsed with 50 mL total volume of lysis buffer by gravity flow at 4°C. Once rinsed, the proteins were eluted from the resin by sequentially incubating the resin with 500 µL of elution buffer containing 50 mM HEPES, 150 mM NaCl, and 200 mM imidazole four times for 5 min at room temperature. Using a nanodrop, the elutions were measured for absorbance at 280 nm to determine if the protein eluted from the resin.

The elutions containing proteins were pooled and concentrated using Amicon Ultra-15, 3 kDa MWCO (Millipore, UFC9003). The concentrated sample was centrifuged for 1 min at 21,000 xg to remove large particles. A Superdex 200 was equilibrated using a buffer containing 50 mM HEPES, 150 mM NaCl, and 1 mM DTT, and the concentrated sample was injected into a 0.5 mL injection loop. The fractions corresponding to TXNL1 were collected, pooled, and concentrated using Amicon Ultra-15, 3 kDa MWCO. Protein samples were flash-frozen using liquid nitrogen for storage at -80°C.

### Purification of TXNL1-deficient proteasomes

*TXNL1* knockout HEK-293T cells were seeded in 50 15 cm plates at 3.5 million cells per plate and grown for 2 days. Cells were then transfected with 10 µg plasmid DNA encoding 2xFLAG-midnolin using PolyJet per plate. The media was replaced 1-day post-transfection and cells were incubated overnight. The cells were then rinsed once with ice-cold PBS before applying 1 mL of buffer containing 40 mM HEPES, 100 mM NaCl, 2mM MgCl_2_, 5mM ATP, 1x HALT protease, 0.5% CHAPS to each 15 cm plate. The cells were collected by scraping and the lysate was incubated at 4°C with end-to-end rotation for 20 min. The lysate was then centrifuged at 21,000 xg for 20 min at 4°C. During the centrifugation, 3.6 mL of anti-FLAG M2 agarose resin (Millipore, A2220) was rinsed four times using lysis buffer. After centrifugation, the supernatant was incubated in a 50 mL conical tube containing the washed anti-FLAG resin with end-to-end rotation for 1.5 h at 4°C. The solution was then passed through a column (BioRad, 7321010) by gravity at 4°C. The conical tube and the column were rinsed with 50 mL of lysis buffer total by gravity flow at 4°C. The column was also rinsed once with 20 mL of size-exclusion chromatography (SEC) buffer containing 40 mM HEPES, 100 mM NaCl, 2mM MgCl, 5mM ATP, 0.05% CHAPS at 4°C.

The resin was then incubated with 1.8 mL of SEC buffer containing 0.5 mg/mL of 3xFLAG peptide (APExBIO, A6001) for 20 min at room temperature. A total of four elutions were performed sequentially and the protein content in each elution was determined by obtaining the absorbance value at 280 nm using a nanodrop. The eluted proteins were pooled and concentrated using an Amicon Ultra-15, 10 kDa MWCO concentrator until 500 µL. The concentrated sample was centrifuged for 1 min at 21,000 xg to remove large particles. A Superose6 column was equilibrated using the SEC buffer for size-exclusion chromatography and the sample was manually injected into a 0.5 mL injection loop. The fractions corresponding to the proteasome were collected, pooled, and concentrated using Amicon Ultra-15, 10 kDa MWCO. Protein samples were flash-frozen using liquid nitrogen for storage at -80°C.

### Binding and degradation assays with purified components

To test TXNL1 binding to proteasomes *in vitro*, 1 µL of each TXNL1 protein (1.7 mg/mL) was diluted in 199 µL of sample buffer and 1 µL of proteasomes (3 mg/mL) were diluted in 99 µL of sample buffer as input. 10 µL of anti-HA beads (Thermo Fisher Scientific, 88836, RRID: AB_2749815) per reaction were washed thrice using 700 µL of CHAPS buffer (100 mM NaCl, 40 mM HEPES pH 7.4, 0.5 % CHAPS, 1x HALT protease/phosphatase inhibitor). After washing, the beads were resuspended in 700 µL of CHAPS buffer containing 4% BSA (40 mg/mL). The beads were blocked for 1 h at 4°C with end-to-end rotation. After blocking, the beads were washed thrice using 700 µL of CHAPS buffer. The beads were resuspended in CHAPS buffer and 10 µL of beads were aliquoted into pre-lubricated Eppendorf tubes (Millipore, CLS3207). HA-TXNL1 protein (15 µg per reaction) was added directly into the beads to a final volume of 500 µL in the CHAPS buffer. The beads were incubated for 1 h at 4°C with end-to-end rotation to immobilize the TXNL1. The beads were then washed thrice using 700 µL of CHAPS buffer and resuspended to 500 µL in CHAPS buffer. 4 µL of proteasomes (3 mg/mL) were added to each reaction. For the chelator experiment, proteasomes were pre-incubated with 6 mM 1,10-phenanthroline (Sigma, 516705) or 1,7-phenanthroline (Sigma, 301841) for 30 min at room temperature before adding the mixture to the TXNL1-bound beads. The beads were then incubated with end-to-end rotation for 1 h at 4°C. The beads were then washed thrice using 700 µL of CHAPS buffer before resuspending the beads in 25 µL of Tris-Glycine sample buffer containing 10% 2-mercaptoethanol. Proteins were eluted from the beads by denaturing at 95°C for 3 min.

To assay for protein degradation, 50 nM purified proteasomes were pre-incubated at room temperature for 30 min with a negative control buffer, 6 mM 1,10-phenanthroline, or 50 µM TXNL1 variants in 100 mM NaCl and 40 mM HEPES pH 7. Then, 50 nM poly-ubiquitinated substrate was added directly to the proteasome mixture along with 5 mM ATP (Sigma, A2383) in a 10 µL reaction volume, incubated for 10 min at 37°C, and quenched with 10 µL of Tris-Glycine sample buffer containing 10% 2-mercaptoethanol. Proteins were denatured at 95°C for 3 min and 8 µL of sample was loaded into 4-20 % Tris-Glycine 15-well pre-cast gels (Thermo Fisher Scientific, XP04205BOX). In-gel fluorescent images were acquired using an Odyssey LI-COR.

### Multiplexed mass spectrometry

Cells were washed twice with 1x PBS, harvested on ice using a cell scraper in 1x PBS, pelleted via centrifugation for 5 min at 1000xg at 4°C, and washed with 1x PBS before resuspension in in 8 M urea (Sigma U5378), 50 mM NaCl, 50 mM EPPS (Sigma E9502) and 1x Protease inhibitor cocktail (Roche 4906845001). After 10 sec of sonication, lysed cells were pelleted, and the protein concentration of clarified sample was determined using a BCA kit (Thermo Fisher Scientific 23225). 100 µg of each sample was incubated for 30 min at 37°C with 5 mM TCEP (Gold Biotechnology) for disulfide bond reduction with subsequent alkylation with 20 mM Iodoacetamide (Sigma I6125) for 20 min at room temperature followed by quenching with 15mM DTT for 15 min under gentle shaking. MeOH-chloroform precipitation of samples was performed as follows: to each sample, 4 parts MeOH was added followed by vortexing, one part chloroform was added followed by vortexing, and finally 3 parts H_2_O was added. After vortexing, the suspension was centrifugated for 5 min at 14000xg and the aqueous phase around the protein precipitate was removed using a loading tip. The precipitate was washed twice with MeOH and resuspended in 200 mM EPPS (pH 8.0) (Sigma E9502) and digested with Trypsin/LysC (Promega V5073) digestion (1:100) at 37°C overnight with gentle shaking.

150 µl of digested samples were labeled by adding 10 µl of TMT reagent (Thermo Scientific A52045, stock: 20 mg/mL in acetonitrile, ACN, Millipore Sigma 34851) together with 50 µl ACN to yield a final ACN concentration of approximately 25% (v/v) for 1 h at room temperature before quenching the reaction with hydroxylamine (Thermo Fisher Scientific 90115) to a final concentration of 0.2% (v/v). The TMTpro-labeled samples were pooled together at a 1:1 ratio, resulting in consistent peptide amount across all channels. Pooled samples were vacuum centrifuged for 1 h at room temperature to remove ACN, followed by reconstitution in 1% FA, desalting using C18 solid-phase extraction (SPE) (200 mg, Sep-Pak, Waters WAT054960), and vacuum centrifugation until near dryness. We fractionated the pooled, labeled peptide sample using BPRP HPLC^32^ and an Agilent 1260 pump equipped with a degasser and a UV detector (set at 220 and 280 nm wavelength). Peptides were subjected to a 50-min linear gradient from 5% to 35% acetonitrile in 10 mM ammonium bicarbonate pH 8 at a flow rate of 0.6 mL/min over an Agilent 300Extend C18 column (3.5 μm particles, 4.6 mm ID and 220 mm in length). The peptide mixture was fractionated into a total of 96 fractions, which were consolidated into 24 super-fractions^33^, of which twelve non-adjacent fractions were analyzed. Samples were subsequently acidified with 1% formic acid (FA, Sigma 94318) and vacuum centrifuged to near dryness. Each super-fraction was desalted via StageTip, dried again via vacuum centrifugation, and reconstituted in 10 µl 5% ACN, 5% FA for LC-MS/MS processing.

Mass spectrometric data were collected on Orbitrap Fusion Lumos instruments coupled to a Proxeon NanoLC-1200 UHPLC. The 100 µm capillary column was packed with 35 cm of Accucore 150 resin (2.6 μm, 150Å; ThermoFisher Scientific) at a flow rate of 340 nL/min. The scan sequence began with an MS1 spectrum (Orbitrap analysis, resolution 60,000, mass range 350-1350 Th, automatic gain control (AGC) target 100%, maximum injection time 118ms). Data were acquired ∼90 min per fraction. The hrMS2 stage consisted of fragmentation by higher energy collisional dissociation (HCD, normalized collision energy 36%) and analysis using the Orbitrap (AGC 200%, maximum injection time 120 ms, isolation window 0.6 Th, resolution 50,000). Data were acquired using the FAIMSpro interface the dispersion voltage (DV) set to 5,000V, the compensation voltages (CVs) were set at -40V, -60V, and -80V, and the TopSpeed parameter was set at 1 sec per CV.

Spectra were converted to mzXML via MSconvert^34^. Database searching included all entries from the mouse UniProt reference Database (downloaded: August 2021). The database was concatenated with one composed of all protein sequences for that database in the reversed order. Searches were performed using a 50-ppm precursor ion tolerance for total protein level profiling. The product ion tolerance was set to 0.03 Da. These wide mass tolerance windows were chosen to maximize sensitivity in conjunction with Comet searches and linear discriminant analysis^35,36^. TMTpro labels on lysine residues and peptide N-termini +304.207 Da), as well as carbamidomethylation of cysteine residues (+57.021 Da) were set as static modifications, while oxidation of methionine residues (+15.995 Da) was set as a variable modification. Peptide-spectrum matches (PSMs) were adjusted to a 1% false discovery rate (FDR)^37,38^. PSM filtering was performed using a linear discriminant analysis, as described previously^36^ and then assembled further to a final protein-level FDR of 1%^37^. Proteins were quantified by summing reporter ion counts across all matching PSMs, also as described previously^39^. Reporter ion intensities were adjusted to correct for the isotopic impurities of the different TMTpro reagents according to manufacturer specifications. The signal-to-noise (S/N) measurements of peptides assigned to each protein were summed and these values were normalized so that the sum of the signal for all proteins in each channel was equivalent to account for equal protein loading. Finally, each protein abundance measurement was scaled, such that the summed signal-to-noise for that protein across all channels equals 100, thereby generating a relative abundance (RA) measurement.

MSstatsTMT^40^ was performed on peptides with > 200 summed SNR across TMT channels. For each protein, the filtered peptide–spectrum match TMTpro raw intensities were summed and log_2_ normalized to calculate protein quantification values (weighted average) and normalized to total TMT channel intensity across all quantified PSMs (adjusted to median total TMT intensity for the TMT channels)^41^. Log_2_ normalized summed protein reporter intensities were compared using a Student’s t-test and p-values were corrected for multiple hypotheses using the Benjamini-Hochberg adjustment^42^.

### RNA Sequencing

HEK-293T cells were grown to 80% confluency in 6-well plates, treated with oxidizing agents (100 µM arsenite and 600 µM hydrogen peroxide) in triplicate for 6 h, and total RNA was isolated using an RNEasy Plus mini kit (QIAGEN, 74134). mRNA-seq libraries were prepared by Innomics and sequenced on a DNBseq platform to obtain 100 bp paired-end reads. Reads were aligned to the GRCh38 primary assembly obtained from gencodegenes.org using qAlign from the QuasR package. Raw counts were filtered to discard transcripts with fewer than 10 total reads in the three untreated *TXNL1* KO replicates, and then the DESeq2 package in R was used to obtain differential expression data and statistics.

## Data Availability

EM maps and models are available under accession numbers EMD-44949, EMD-44952, and PDB 9BW4. The mass spectrometry proteomics data are available via ProteomeXchange with identifier PXD052933. RNA sequencing data are available through GEO under the accession number GSE271951.

